# A Divergent Cytochrome c in Malaria Parasites with an Anomalously Low Redox Potential

**DOI:** 10.64898/2025.12.23.696199

**Authors:** Kade M. Loveridge, Otessa D. Olsen, Samuel R. Scherer, Tanya J. Espino-Sanchez, Henry Wienkers, Christopher P. Hill, Michael S. Kay, Paul A. Sigala

**Author notes:** These authors contributed equally to this work (Kade M. Loveridge, Otessa D. Olsen).

## Abstract

Eukaryotic cytochrome (cyt) *c* is a highly conserved mitochondrial protein central to cellular respiration, featuring a covalently attached hexacoordinate heme whose redox potential is tuned by axial His/Met ligands and surrounding residues to support electron transport chain (ETC) function. We have identified an unrecognized lineage of eukaryotic cyt *c* homologs in Apicomplexa, a phylum of intracellular pathogens that includes *Plasmodium falciparum* malaria parasites. *P. falciparum* cyt *c*-2 (Pfcyt *c*-2) exemplifies this divergent lineage and has an unusual pentacoordinate heme despite conservation of His/Met ligands. We determined that Pfcyt *c*-2 has a redox potential of –278 mV that is over 500 mV lower than canonical cyt *c* homologs (+250 mV) and contradicts a conserved ETC role. This anomalous redox potential is lower than any natural monoheme *c*-type cyt. Nevertheless, Pfcyt *c*-2 displays canonical thermostability and low-level peroxidase activity, while showing signs of elevated structural heterogeneity. These results reveal a new clade of eukaryotic cyt *c* variants with divergent biochemical properties and biological roles, opening new scaffolds for mechanistic discovery and redox engineering.

## RESULTS

*C*-Type cytochromes (cyt) are defined by covalent heme attachment through their conserved CXXCH motif but span diverse structural classes^1, 2^. Eukaryotic mitochondrial cyt *c* represents the archetypal Class I *c*-type cyt^3^. Its hexacoordinate His/Met ligation enforces a low-spin heme with minimal electronic rearrangement between oxidation states, yielding a redox potential of ∼250 mV that enables electron transfer between Complex III and Complex IV in the mitochondrial ETC^4, 5^. Although cyt *c* also functions as a peroxidase in animals during cellular apoptosis, these roles are secondary to its role as an electron shuttle. Compared with other heme proteins, such as cytochrome P450s that catalyze diverse oxidative reactions, cyt *c* proteins are generally viewed as chemically conservative scaffolds^6, 7^.

*Plasmodium falciparum*, the causative parasite of severe malaria, encodes two cyt *c* paralogs, a canonical Pfcyt *c* (Uniprot Q8IM53) with conserved ETC function, and a highly divergent Pfcyt *c*-2 (Uniprot Q8I6T6) that features extensive changes in the heme crevice residues proximal to the conserved axial Met (Met81, using bovine numbering)^8^. Pfcyt *c*-2 is hemylated on its CXXCH motif by the parasite holocytochrome c synthase (HCCS) and trafficked to mitochondria^8, 9^. Despite conservation of axial His/Met ligands, Pfcyt *c*-2 displays spectroscopic features (by UV-vis and electron paramagnetic resonance) indicative of a pentacoordinate, high-spin heme in the oxidized state that is able to bind external ligands, including imidazole and H_2_O_2_^8^. In the reduced state, however, UV–vis spectra of Pfcyt *c*-2 resemble those of a low-spin hexacoordinate heme with undefined axial ligands^8^.

These observations and X-ray structural studies, which revealed preferential crystallization of a domain-swapped dimer (DSD) over the monomeric form that predominates in solution, suggest that structural heterogeneity in the heme crevice region destabilizes axial coordination by Met81^8^. The Pfcyt *c*-2 DSD also demonstrated covalent heme attachment, proximal His19 ligation, and retention of secondary structure elements typical of cyt *c* proteins (Fig. S1, PDB: 7TXE)^8^. These biochemical features of Pfcyt *c*-2 are highly unusual for a eukaryotic cyt *c* and suggest that it represents a distinct branch of cyt *c* variants with divergent cellular functions.

To trace the evolutionary origins of the parasite cyt *c* paralogs, we queried the canonical Pfcyt *c* and divergent Pfcyt *c*-2 across all major eukaryotic lineages. Cyt *c*-2 homologs were detected only for intracellular parasites in the phylum Apicomplexa and their closest free-living relative, *Vitrella brassicaformis*, suggesting that this molecular divergence accompanied the evolutionary appearance of parasitism^10^. Comparative analyses also revealed an unrecognized clade, which we term cyt *c*-1.5, that branches between cyt *c* and *c*-2 (Fig. S2). Sequence similarity networks resolved three distinct clusters (*c, c*-1.5, *c*-2), and phylogenetic reconstruction indicates that cyt *c*-1.5 and *c*-2 arose from two sequential duplication events (Fig. 1A,B)^11-13^. Many apicomplexan lineages, including *Plasmodium* parasites, appear to have lost cyt *c*-1.5 and retained only cyt *c*-2.

**Figure 1.**
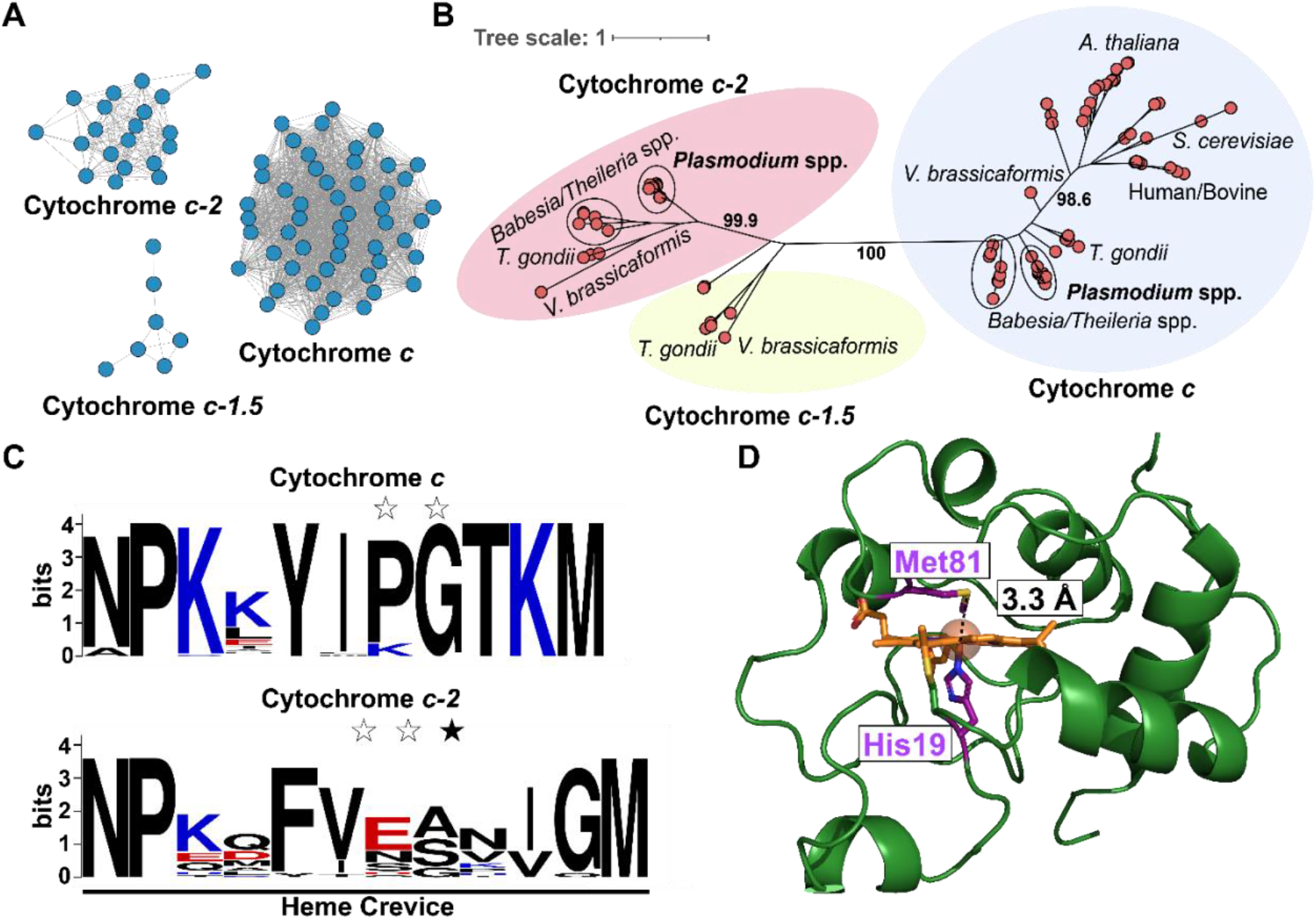
Bioinformatic analysis of cyt *c*-2 homologs. Sequences of cyt *c, c*-1.5, and *c*-2 were analyzed by (A) sequence similarity network (54% identity threshold), (B) phylogenetic tree, and (C) heme crevice sequence logo. Blue and red depict positively and negatively charged residues, respectively. White stars indicate loss of structural proline/glycine, and the black star marks an inserted residue. (D) Model of Pfcyt *c*-2 bound to heme *c* with labeled S-Fe distance (N and C-termini truncated, full model provided in Fig. S5).

Multiple sequence alignments highlight insertions between α helices α1 and α2 (Fig. S2). The heme crevice is the most conserved region of cyt *c* proteins and shows striking variability in cyt *c*-2 homologs, including charge reversals, disruption of structural residues, and an insertion (Fig. 1C)^14, 15^. Multiple residues that stabilize Met81 coordination have diverged, consistent with the oxidized pentacoordinate state of Pfcyt *c*-2. Despite alterations in the heme crevice region of Pfcyt *c*-2, hallmark features of canonical cyt *c* remain intact, including Met81 and residues required for folding, HCCS recognition, and porphyrin-ring contacts (Fig. S3)^3, 15^.

Because a monomeric crystal structure was not obtainable, we modeled 16 unique cyt *c*-2 proteins with heme *c* using AlphaFold3^16^ (Fig. S4)^17^. The majority of cyt *c*-2 models maintained a general cyt *c* fold but lacked stable Met–Fe coordination and were predicted in one of three states: (i) Met81 proximal but unbound, (ii) Met81 partially displaced, or (iii) Met81 outside/adjacent to the heme crevice (Fig. S4 and S5). Pfcyt *c*-2 was predicted to adopt state (i), in which Met81 sulfur lies 3.3 Å from the heme iron with its sulfur lone-pair electrons rotated away from the metal center and thus mispositioned to coordinate iron (Fig. 1D).

Perturbations to axial heme coordination and the local active site environment of Pfcyt *c*-2 suggested alterations to its redox properties. Using comparative dye-based reduction by UV– vis absorbance^18^, we first determined midpoint potentials of 237 ± 12 mV and 216 ± 3 mV for bovine cyt *c* and Pfcyt *c*, which were similar to other canonical homologs and consistent with their roles in the ETC (Fig. S6)^19-21^. In contrast, Pfcyt *c*-2 was not reducible under the same conditions (Fig. S7), prompting us to test a panel of reductants (Fig. S8). Pfcyt *c*-2 could only be reduced by sodium dithionite (E° ≈ –660 mV) but not by other reductants of higher redox potential (Fig. S8), indicating an exceptionally low redox potential. Guided by this observation, we measured a midpoint potential of –278 ± 6 mV for Pfcyt *c*-2, a value independently confirmed using two distinct dyes (Fig. 2, Fig. S9) ^18^.

**Figure 2.**
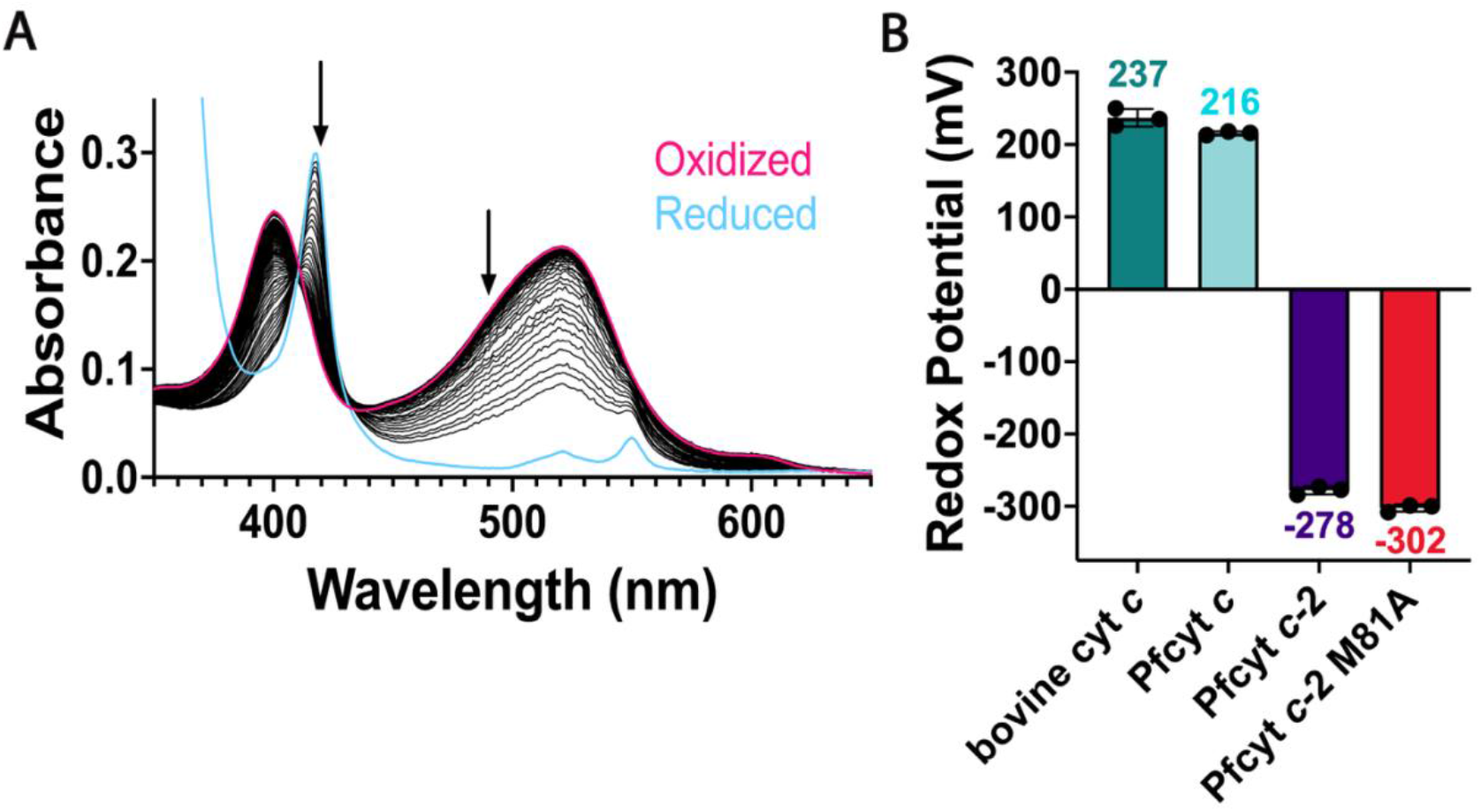
Determination of the redox potential of Pfcyt *c*-2. (A) Representative UV–vis spectra of the oxidized (pink) to reduced (blue) transition during equilibrium titration of Pfcyt *c*-2 and Safranin dye. Arrows mark wavelengths (420, 490 nM) used to calculate oxidized/reduced fractions for Nernst analysis. (B) Midpoint redox potentials of bovine cyt *c*, Pfcyt *c*, and Pfcyt *c*-2 shown as the mean of three independent determinations ± SD (versus standard hydrogen electrode).

This anomalously low redox potential represents a >500 mV decrease relative to canonical cyt *c*. Such a low potential is incompatible with ETC electron shuttling and lies far below the previously lowest reported monoheme cyt, *Geobacter sulfurreducens* PccH (–24 mV)^22^. To our knowledge, Pfcyt *c*-2 has the lowest redox potential described for any naturally occurring monoheme *c*-type cytochrome^22-24^. Comparable values have only been achieved in artificially engineered cyt *c* M81C mutants, where axial replacement with Cys introduces an electron-donating thiolate ligand that stabilizes the Fe(III) state and drives the potential downward, or in multiheme cytochromes, where electronic communication between closely packed heme centers redistributes charge and lowers the midpoint potential through cooperative redox coupling^25, 26^.

Given the high sequence divergence of Pfcyt *c*-2 and the evidence that the distal Met81 does not coordinate heme in the oxidized state, we asked whether this residue might still contribute to the low redox potential as a basis for its conservation. An M81A Pfcyt *c*-2 mutant was efficiently recognized and hemylated by human HCCS (Fig. S10)^27^. UV–vis spectra of the oxidized and reduced forms of M81A showed only subtle differences, including mild broadening of the Soret peak, and the oxidized spectra of WT and M81A in the presence of imidazole were nearly superimposable (Fig. S11). The mutant displayed a small but reproducible ∼20 mV decrease in redox potential, in contrast to the >250 mV decrease previously observed for the corresponding mutant in human cyt *c*^27^. This modest decrease in Pfcyt *c*-2 strongly suggests that Met81 likely modulates the heme environment indirectly (Fig. 2B, Fig. S12). These findings argue against a role for the conserved Met81 as an axial ligand in the reduced state but instead suggest that it contributes structurally to shaping the electronic properties of the heme crevice. This suggestion is consistent with AlphaFold predictions, which place Met81 within or adjacent to that region but misposition it to coordinate the heme iron.

The physical features of Pfcyt *c*-2 that underpin this unusually low reduction potential remain to be elucidated, but loss of stable Met81 coordination alone is only expected to reduce the redox potential ∼250 mV based on mutagenesis studies of canonical cyt *c* proteins ^27-30^. AlphaFold models of cyt *c*-2 homologs suggest a shortening of the His19-Fe bond by ∼0.1 Å, a feature also present in the DSD Pfcyt *c*-2 dimer, which would stabilize the Fe(III) state (Fig. S13) and may contribute to lowering the redox potential^14, 31, 32^. Continuum electrostatics calculations also suggest enhanced (≥twofold) negative charge within 10 Å of the heme relative to canonical cyts *c*, a shift predicted to further lower the E°′ (Fig. S14)^33-35^. These and other factors may contribute to the >500 mV decrease in redox potential and can be dissected in future structural and computational studies.

In mammals, cyt *c* acquires pro-apoptotic functions when Met81 is displaced by proteins or lipids such as cardiolipin^3, 6^. This disruption of the native Met ligation exposes the heme to unlock low-level peroxidase activity, a property also reproduced in synthetic pentacoordinate cyt *c* mutants^36^. Because Pfcyt *c*-2 is natively pentacoordinate, we predicted that it might display enhanced peroxidase activity. To test this prediction, we performed kinetic assays with H_2_O_2_ and guaiacol that revealed that bovine cyt *c*, Pfcyt *c*, and Pfcyt c-2 all had very similar *k*_cat_ values of 0.17 s^−1^, 0.07 s^−1^ and 0.10 s^−1^, respectively (Fig. 3A, Fig. S15). Thus, despite evidence for constitutively exposed and pentacoordinate heme, Pfcyt *c*-2 retains basal peroxidase activity typical of higher eukaryotic cyt *c* proteins^37^. Enhanced active-site accessibility and pentacoordinate heme are therefore insufficient to enhance peroxidase function beyond that expected for a cyt *c* with axial His coordination^38, 39^.

**Fig. 3.**
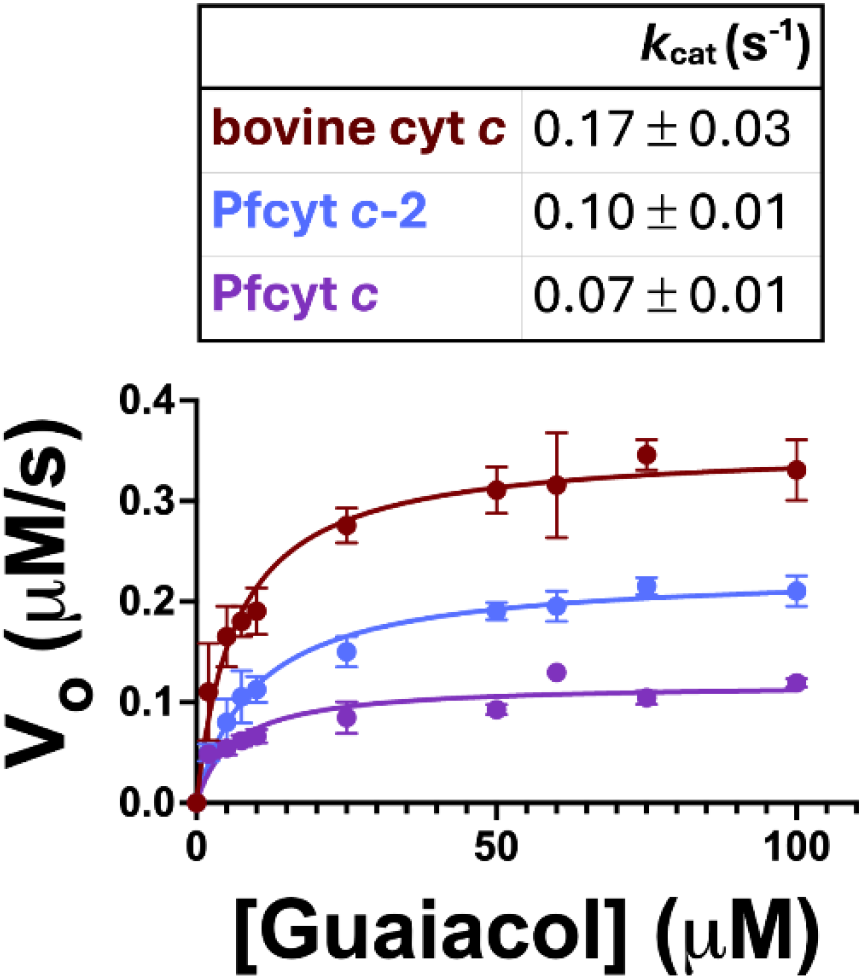
Determination of Pfcyt *c*-2 peroxidase activity and stability. Michaelis-Menton plot of peroxidase activity for bovine cyt *c*, Pfcyt *c*, and Pfcyt *c-*2 with varying guaiacol concentrations. Each point is the mean V_o_ of three independent replicates. Inset shows mean *k*_cat_ values determined from assays at three different enzyme concentrations (Fig. S15).

The extensive sequence insertions in Pfcyt *c*-2 and the inability to crystallize a physiological monomer raised the question of whether the protein might be destabilized relative to typical cyt *c* proteins. Thermal melt analysis by circular dichroism revealed a temperature-dependent unfolding transition in Pfcyt *c*-2 with a midpoint T_m_ of 75 °C, only slightly below the 82 °C observed for bovine cyt *c*, indicating that the Pfcyt *c*-2 fold remains thermostable (Fig. S16) despite evidence for local static or dynamic structural disorder in the heme crevice region^8, 27^.

Pfcyt *c*-2 has evolved sequence, structural, and biochemical features that are highly unusual for a mitochondrial cyt *c*, including loss of axial coordination by Met81, which leads to pentacoordinate heme with axial His ligation, redox spin-state switching, and an anomalously low reduction potential approaching -300 mV. We propose that these properties are distinguishing features of this newly identified cyt *c*-2 lineage and are likely general to other apicomplexan parasites, including the human pathogens *Toxoplasma gondii* and *Babesia* spp. Results in this and our prior study argue against a model that enhanced peroxidase activity or roles in oxidative protection underpin the cellular function of Pfcyt *c*-2^8^. Instead, the open heme pocket and strongly shifted redox chemistry point toward alternative functions, potentially binding or chemically transforming small molecules or gases in a signaling or catalytic capacity^7, 24^. The retention of Pfcyt *c*-2 in *Plasmodium* parasites, which have otherwise substantially reduced their heme proteome, underscores durable evolutionary pressure to preserve the atypical sequence and functional properties of this bioinorganic scaffold and its contribution to parasite mitochondrial metabolism^8, 40, 41^. It remains a key future challenge to elucidate the divergent cellular function of Pfcyt *c*-2 and related homologs.

Although canonical cyt *c* proteins are specialized for electron transfer, directed-evolution studies demonstrate that this scaffold can be reprogrammed for abiological catalysis. Mutations supporting C-Si bond-forming activity disrupt the native Met–Fe ligation, generating a pentacoordinate heme with increased solvent accessibility and anionic charge at the distal face^42-44^. Notably, Pfcyt *c*-2 naturally exhibits several of these same features, which may similarly impart a predisposition toward noncanonical promiscuous reactivity that may be further refined through protein engineering to perform novel functionalities in synthetic biology and/or industrial chemistry. More broadly, properties of Pfcyt *c*-2 can guide ongoing efforts to understand and engineer novel heme protein functions.

## SUPPORTING INFORMATION

Includes Materials and Methods, AlphaFold3 models and analyses, UV-vis spectra and Nernst plots for all dyes and cytochromes, Michaelis-Menton curves, and circular dichroism (CD) data (PDF).

## ACKNOWLEDGEMENTS

We thank members of the Sigala lab for helpful discussions. This work was supported by NIH grants R35GM133764 and R21AI185746 (P.A.S.) and the CHEETAH Center Grant U54AI170856 (M. S. K.). K.M.L. was supported by a predoctoral fellowship from the American Heart Association. DNA synthesis and sequencing were performed using core facilities at the University of Utah.

