## Supporting Information for "A Divergent Cytochrome c in Malaria Parasites with an Anomalously Low Redox Potential"

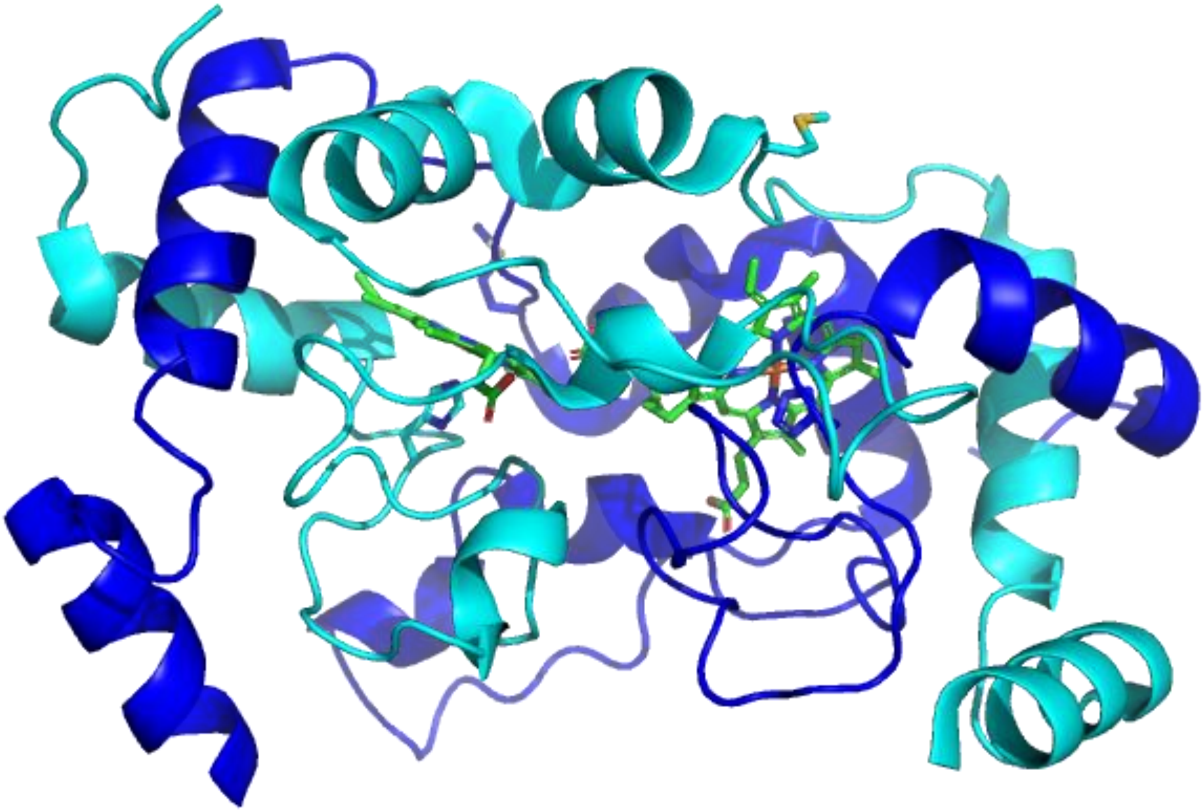

**Figure S1. Crystal structure of the Pfcyt *c*-2 domain-swapped dimer (DSD).** The two Pfcyt *c*-2 monomers are shown in blue and cyan. Heme cofactors are displayed in green (PDB: 7TXE)[1].

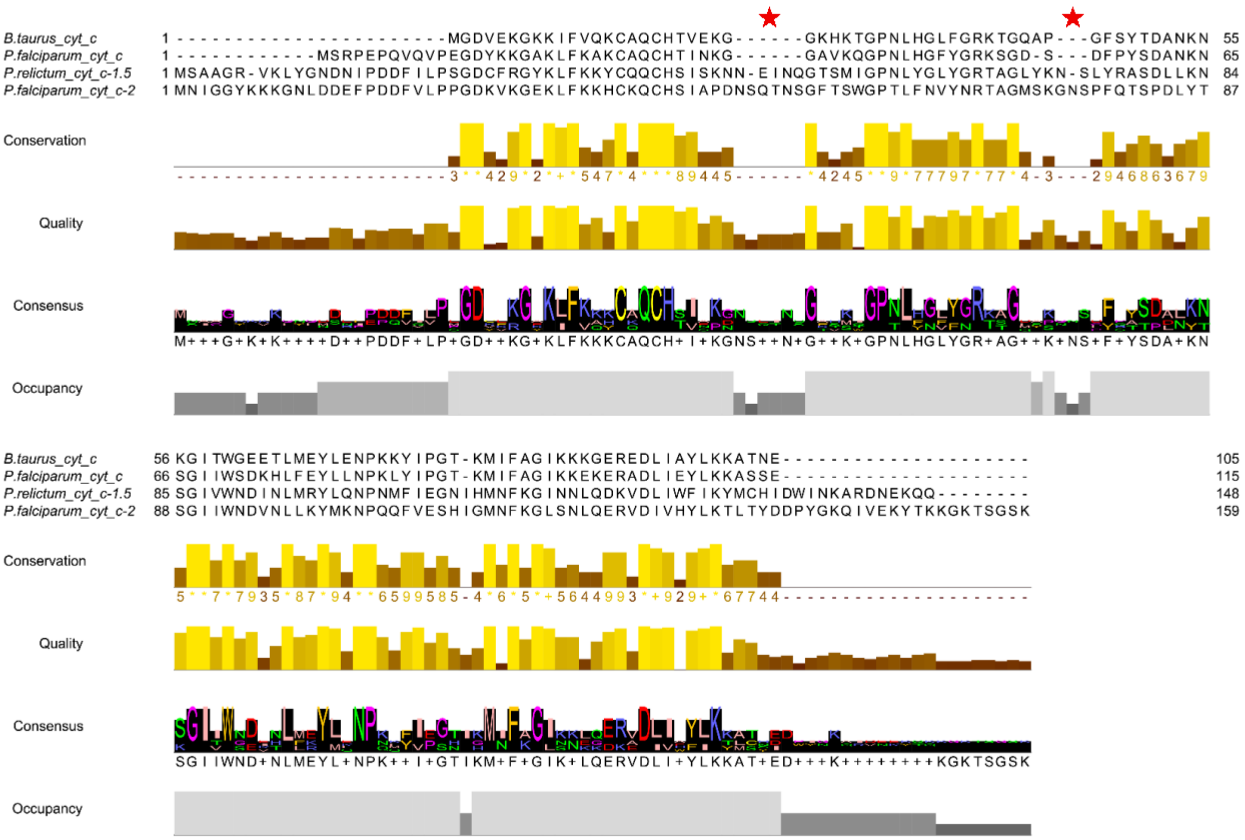


**Figure S2. Sequence alignment of representative cytochromes belonging to the *c*, *c*-1.5, or *c*-2 lineages**. Sequences were aligned in Clustal Omega and visualized with Jalview. Red stars indicate two insertions that occur between α-helices 1 and 2.

```
                                                                 CXXCH
P. falciparum cyt c-2      MNIGGYKKKGNLDDEFPDDFVLPPGDKVKGEKLFKKHCKQCHSIAPDNSQTNSGFTSWGP     60
Bovine cyt c               -----------------------MGDVEKGKKIFVQKCAQCHTVEKG------GKHKTGP     31
P. falciparum cyt c        ------------MSRPEPQVQVPEGDYKKGAKLFKAKCAQCHTINKG------GAVKQGP     42
                                                        G7   F11                      P31
                                                                       K73    T79 F83
P. falciparum cyt c-2      TLFNVYNRTAGMSKGNSPFQTSPDLYTSGIIWNDVNLLKYMKNPQQFVESHIGMNFKGLS     120
Bovine cyt c               NLHGLFGRKTGQAP---GFSYTDANKNKGITWGEETLMEYLENPKKYIPG-TKMIFAGIK     87
P. falciparum cyt c        NLHGFYGRKSGD-S---DFPYSDANKNSGIIWSDKHLFEYLLNPKLYIPG-TKMIFAGIK     97
                                                   N53    W60     Y68 P72  P77  M81
                                      Y98
P. falciparum cyt c-2      NLQERVDIVHYLKTLTYDDPYGKQIVEKYTKKGKTSGSK     159
Bovine cyt c               KKGEREDLIAYLKKATNE---------------------     105
P. falciparum cyt c        KEKERADLIEYLKKASSE---------------------     115
                                  L95
```

| Residue | Functional Role[2, 3] |
|---|---|
| G7, F11, L95, Y98 | N and C termini residues that interact and are crucial for protein folding and stability. F11 is important for hemylation as it is recognized by HCCS. |
| C15, C18, H19 (CXXCH) | C15 and C18 are sites of heme attachment. H19 is the proximal axial ligand of heme iron. Together they make up the conserved CXXCH motif diagnostic of *c*-type cytochromes. Interestingly, Pfcyt *c*-2 does have an uncommon K in the second position of the CXXCH motif. |
| P31 | Structural proline which is important for positioning of H19 for heme iron coordination. |
| N53, T79 | Part of hydrogen bonding network with Y68 and M81 that modulates redox potential and stabilizes M81 coordination. |
| W60 | Bulky, hydrophobic, and aromatic nature contributes to global protein stability and modulating electrostatic environment around heme. |
| Y68 | Participates in hydrogen bonding network with N53, T79, M81 that is important for redox potential and M81 axial coordination. |
| P72 | Structural proline that shield heme from solvent and positions M81 for stable coordination. |
| K73 | Positively charged residue near partially exposed heme center that is important for protein interactions and apoptotic activity in metazoans. |
| P77 | With G78 makes β turn that positions Met81 for axial coordination |
| M81 | Sixth axial ligand for canonical cytochromes *c* |
| F83 | Important for interactions with partner redox proteins and electron transfer. |

**Figure S3. Pfcyt *c*-2 retention of highly conserved mitochondrial cyt *c* residues.** Sequence alignment of Pfcyt *c*-2, Pfcyt *c*, and bovine cyt *c* highlighting residues that are highly conserved across canonical cyt *c* proteins. Residues are numbered based on the bovine numbering system. The table describes the role each amino acid is proposed to perform. Magenta indicates substitution of key residues in Pfcyt *c*-2; green indicates conservation.

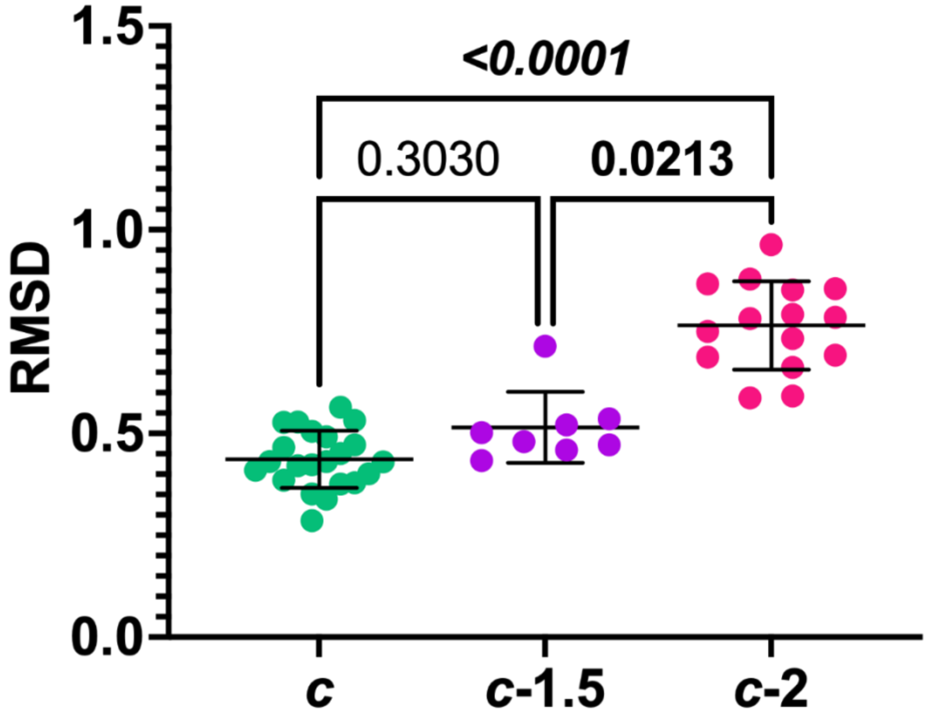


**Figure S4. Structural comparison of experimental and AlphaFold-predicted cyt *c* homologs.** Structural models for the sequences used in Figure 1 include canonical cyts *c* (3 crystal structures[4, 5], 20 models), cyts *c*-1.5 (8 models), and cyts *c*-2 (16 models). These structures were aligned to bovine cytochrome *c* (PDB: 2B4Z)[6] using the PyMOL align command to calculate Cα RMSD values. Statistical differences among groups were assessed using a Kruskal–Wallis test followed by Dunn's post hoc test. The p-values are shown.

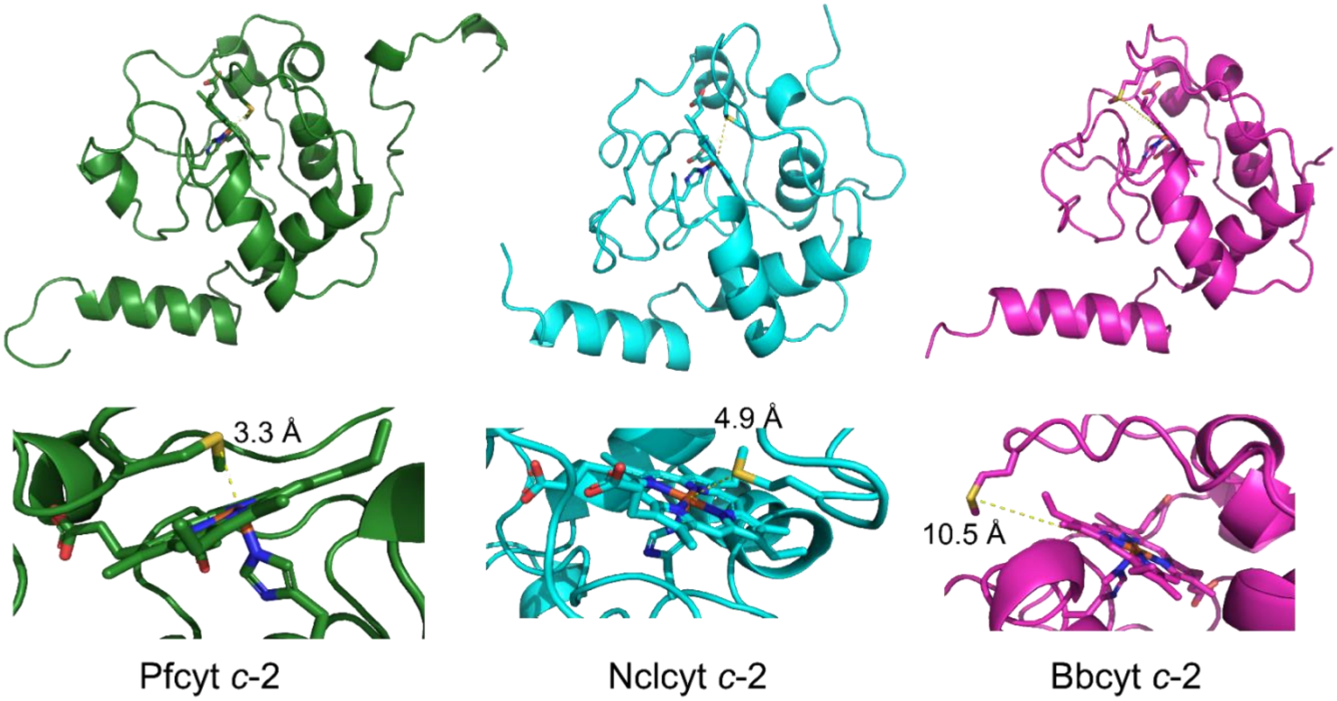


**Figure S5. Predicted structural variants of cyt *c*-2 homologs.** (Top) Representative AlphaFold3-derived cyt *c*-2 models from *Plasmodium falciparum*, *Neospora caninum* (Uniprot: F0VHN6), and *Besnoitia besnoiti* (Uniprot: A0A2A9MB16) are shown. (Bottom) Zoomed in view of the heme crevice region in each homolog. The S–Fe distance of Met81 is annotated to highlight differences in distal ligand positioning relative to the heme iron.

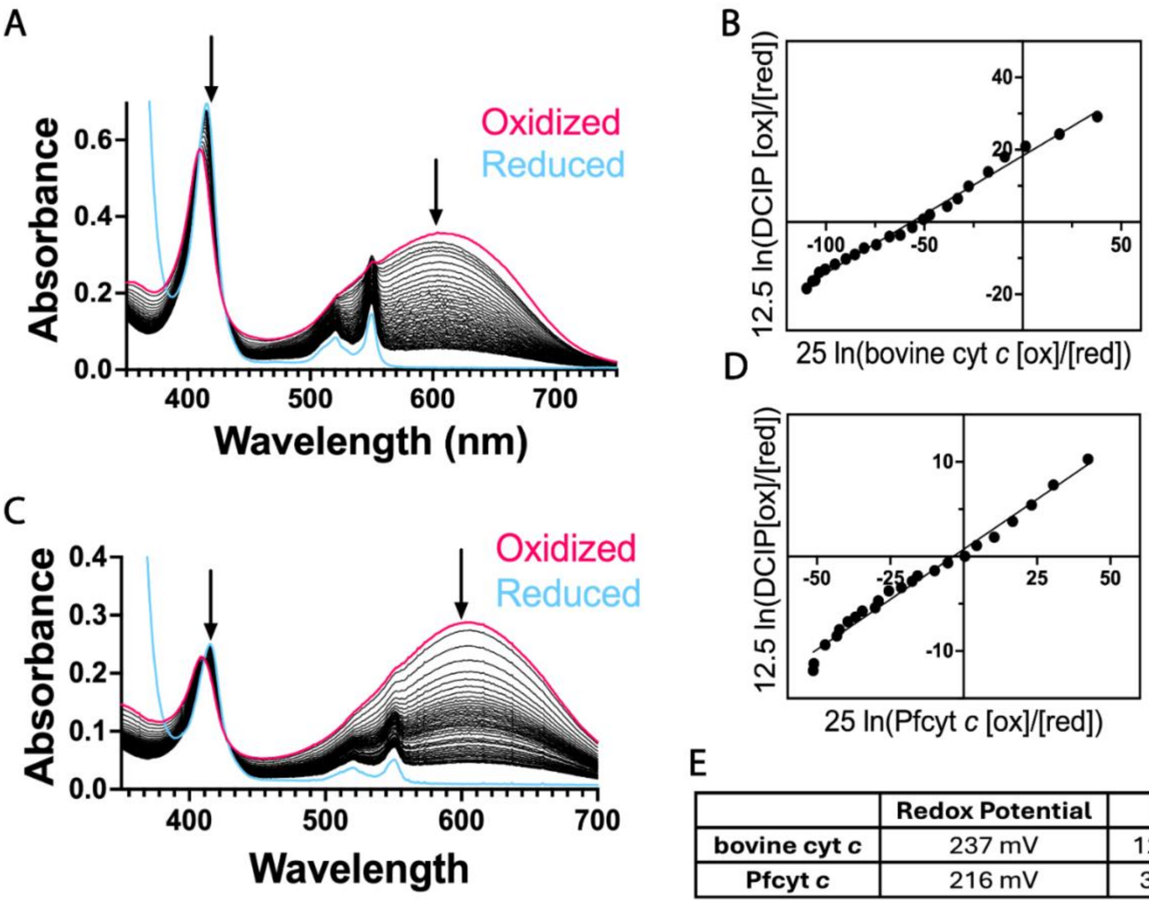


E

| | Redox Potential | SD |
|---|---|---|
| **bovine cyt *c*** | 237 mV | 12 mV |
| **Pfcyt *c*** | 216 mV | 3 mV |

**Figure S6. Redox potential determination of Btcyt *c* and Pfcyt *c*.** Representative UV-vis spectra for bovine cyt *c* (A) and Pfcyt *c* (C) with DCIP dye. Arrows indicate wavelengths (420 and 602 nm) used to determine the ratio between oxidized and reduced species. Nernst plots showing linear relationship between DCIP and Btcyt *c* (B) and Pfcyt *c* (D). (E) Shows the redox potential versus the standard hydrogen electrode as an average of three replicates with standard deviation.

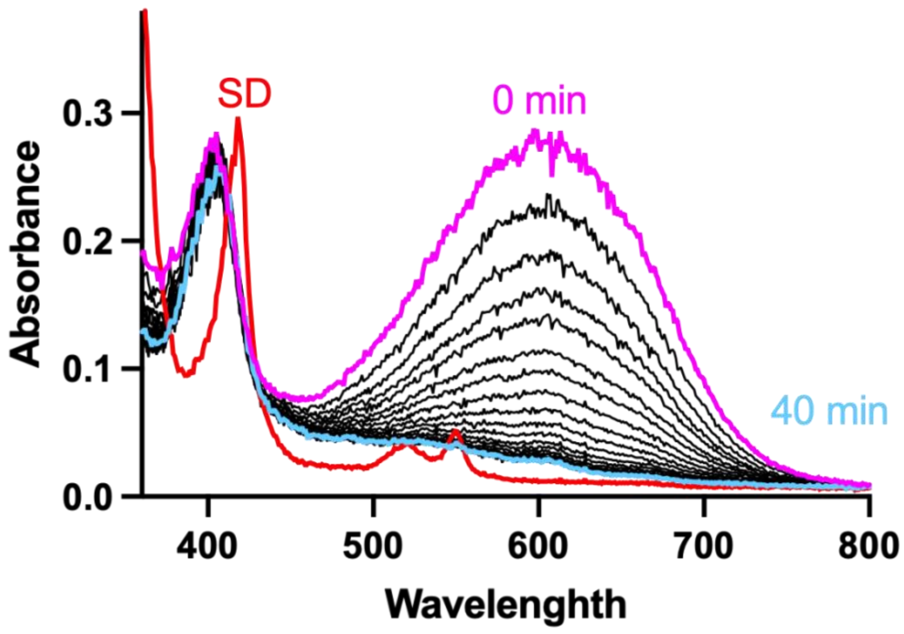


**Figure S7. UV–vis spectra of Pfcyt *c*-2 and DCIP.** Representative UV-vis traces showing the selective reduction of DCIP (E°=217 mV) over 40 minutes in the assay mixture. After 40 minutes, sodium dithionite (SD) to fully reduce DCIP and Pfcyt *c*-2.

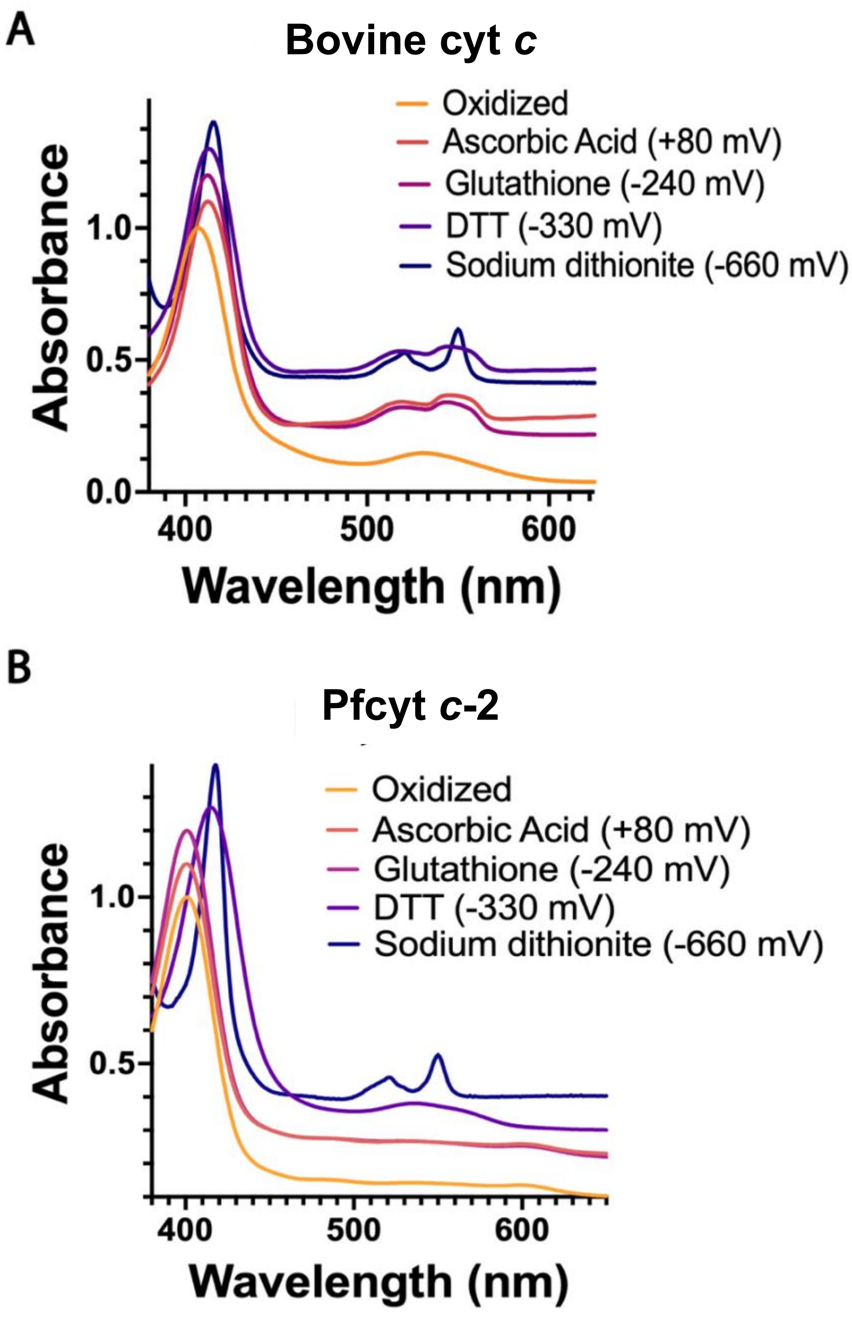


**Figure S8. UV-vis spectra of bovine cyt *c* and Pfcyt *c*-2 with reducing agents.** Absorbance values were normalized to the Soret peak and shifted vertically to minimize overlap. (A) Bovine cyt *c* is reduced by all the reducing agents screened as indicated by the emergence of α,β- peaks at ~ 520 and 550 nm. (B) PfCyt *c*-2 was not reduced with ascorbic acid or glutathione. DTT treatment elicited a broadened, red-shifted Soret peak accompanied by the emergence of a broad ~530 nm feature, consistent with the low-spin, hexacoordinate species observed when adding exogenous ligans such as imidazole or $H_2O_2$[1], in contrast to the sharp red-shifted Soret band and well-resolved α/β bands characteristic of reduced Pfcyt *c*-2.

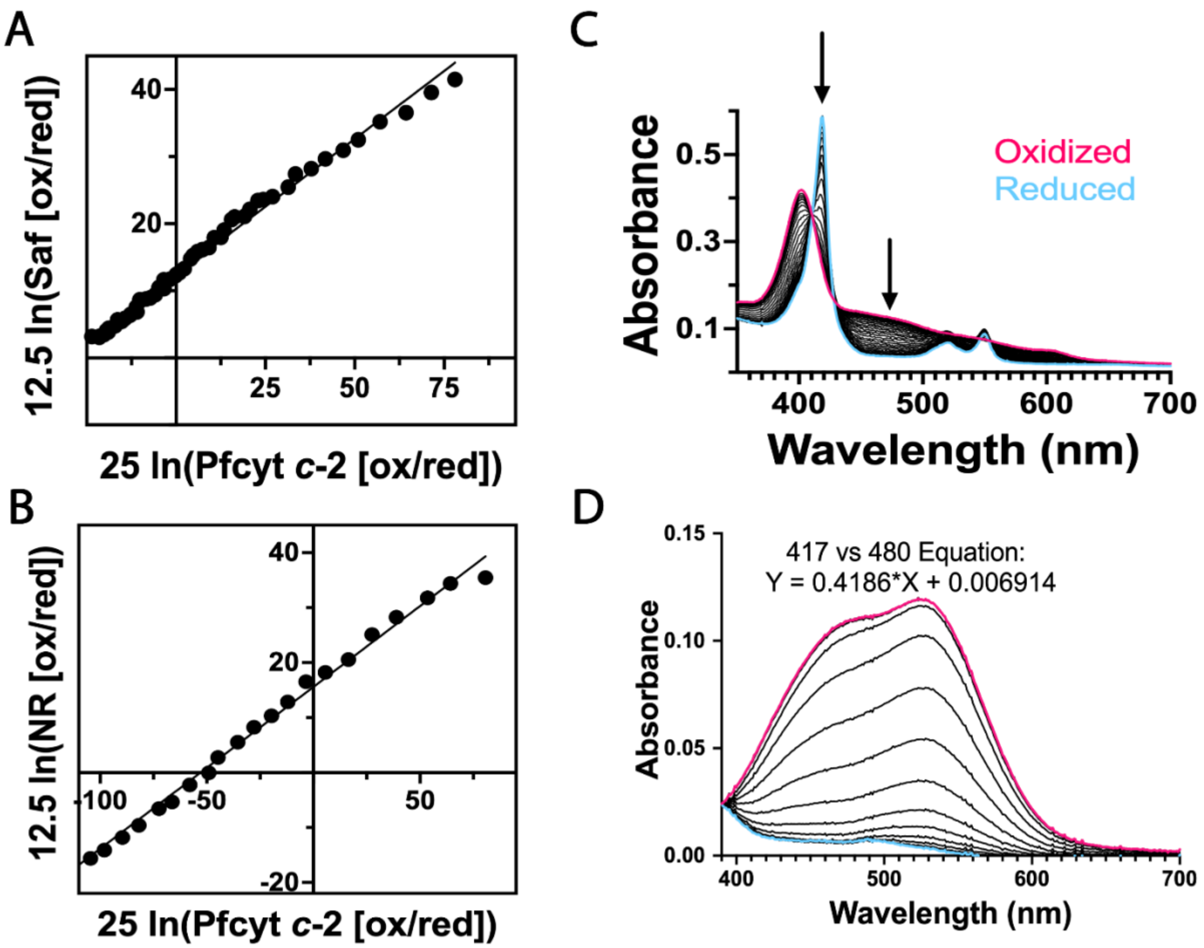


**Figure S9. Additional data for redox potential determination of Pfcyt *c*-2**. Representative Nernst plots showing linear relationship between Pfcyt *c*-2 and (A) Safranine T (-289 mV) and (B) Neutral Red (-325 mV) used to determine the redox potential of Pfcyt *c*-2 to be -278 ± 6 mV and -307 ± 6 mV. (C) Representative UV-vis spectra for Pfcyt *c*-2 with Neutral Red with arrows indicating wavelengths (417 and 480 nm) used to determine the ratio between oxidized and reduced species. (D) UV–vis spectra of Neutral Red undergoing reduction alone, which were used to derive a regression-based correction function describing the relationship between absorbance at 480 nm and 417 nm. The derived 417/480 function was applied during the analysis of each Pfcyt *c*-2 titration to estimate and subtract the Neutral Red contribution to the Soret region caused by spectral overlap.

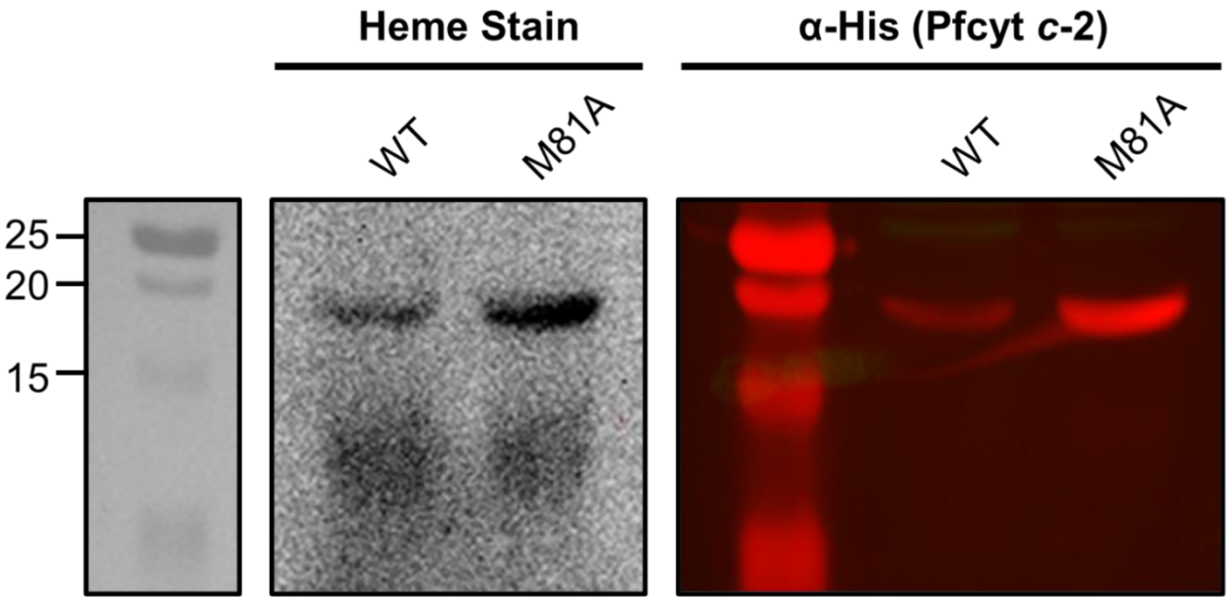


**Figure S10. Heme stain of WT and M81A Pfcyt *c*-2.** WT and M81A Pfcyt *c*-2 were expressed in bacteria as N-terminal fusions with 6xHis (co-expressed with human HCCS). Lysates were fractionated by SDS-PAGE and transferred to a nitrocellulose membrane. Covalent heme attachment was detected using chemiluminescent heme stain. WT and M81A expression were determined using α-His antibodies and western blot analysis.

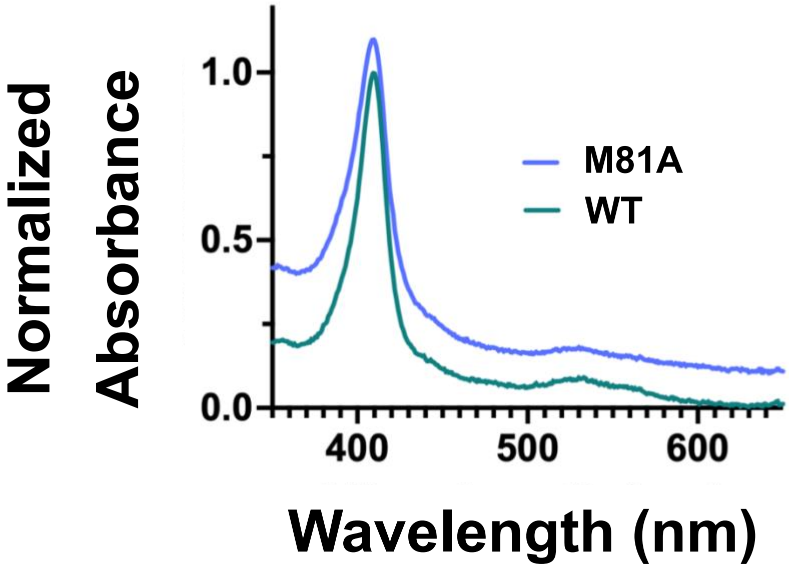


**Figure S11. UV-vis spectra for WT and M81A Pfcyt *c*-2 in the presence of 1 mM imidazole.** Absorbance values were normalized to the Soret peak and offset vertically.

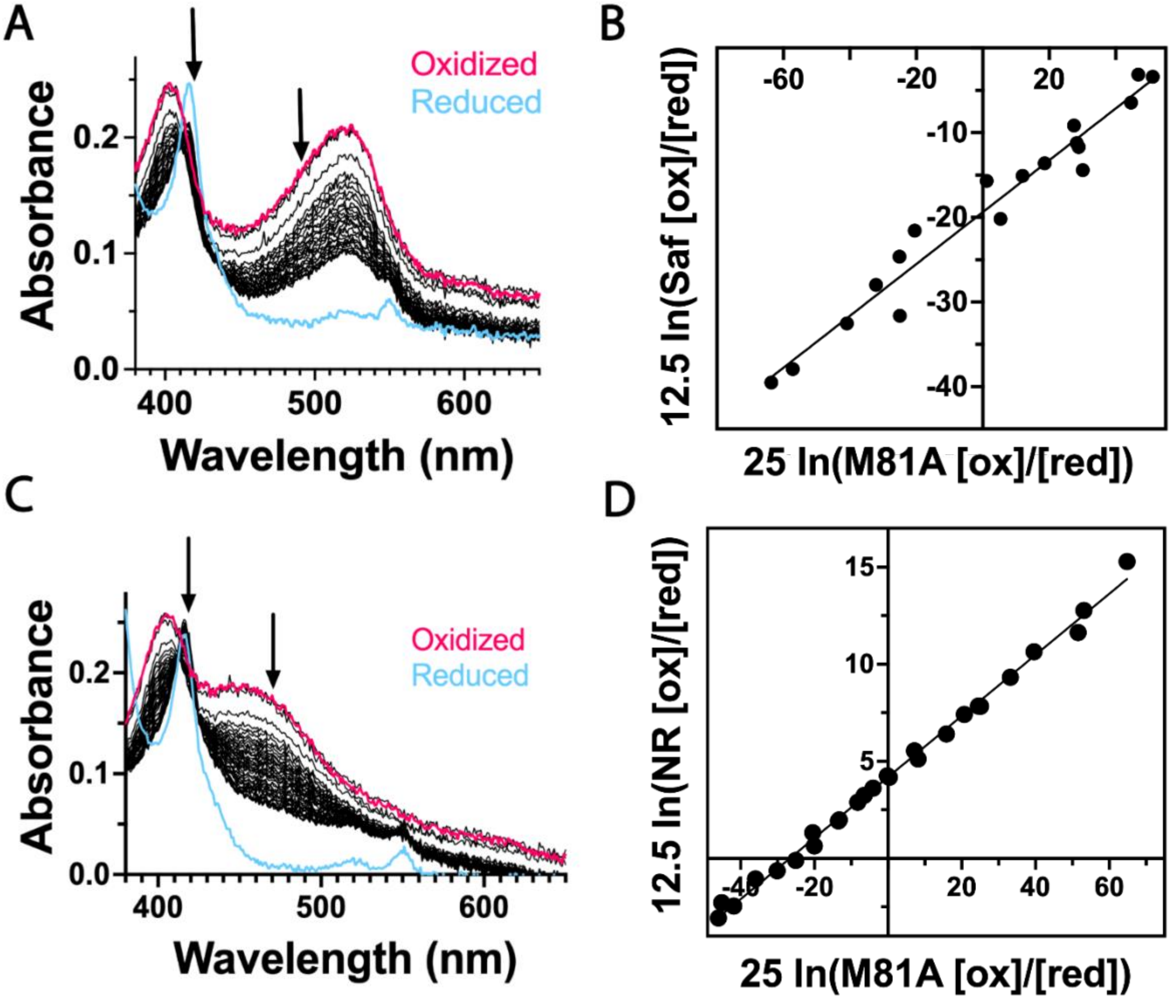


**Figure S12. Determination of M81A Pfcyt *c*-2 redox potential.** (A) and (B) show the UV-vis spectra and Nernst plots respectively for M81A with Safranine T, yielding a potential of -302 ± 5 mV. (C) and (D) show the UV-vis spectra and Nernst plots respectively for M81A with Neutral Red yielding a potential of -322 ± 7 mV.

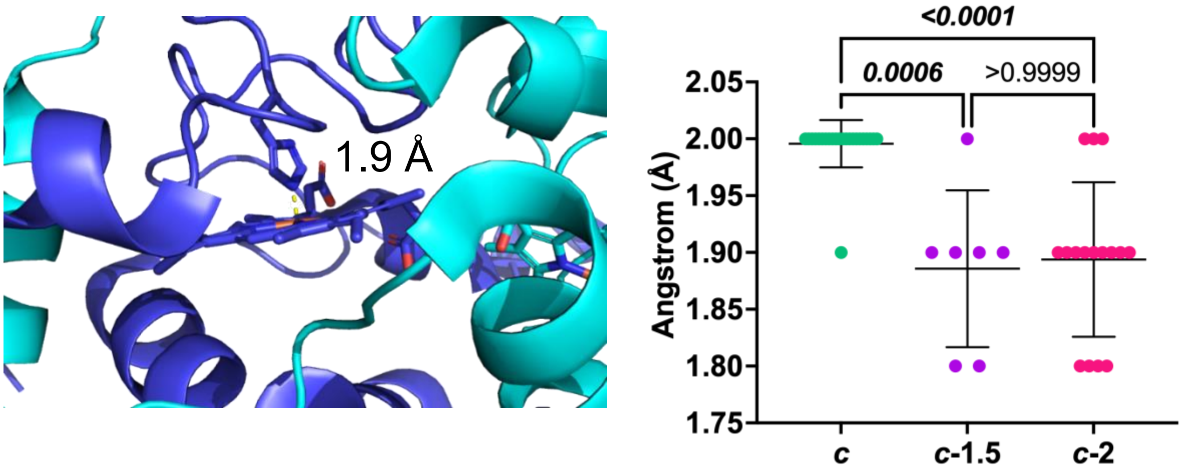


**Figure S13. Measurement of His19-Fe bond distance of Pfcyt *c*-2 DSD and predicted cyt *c*-2 homologs.** Left: Structure of the Pfcyt *c*-2 DSD (7TXE) with annotated length of the His-Fe bond distance. Right: Measured His-Fe bond distance of solved crystal structures or AlphaFold3 predictions from Fig. S3. Statistical differences among groups were assessed using a Kruskal–Wallis test followed by Dunn's post hoc test. The p-values are shown.

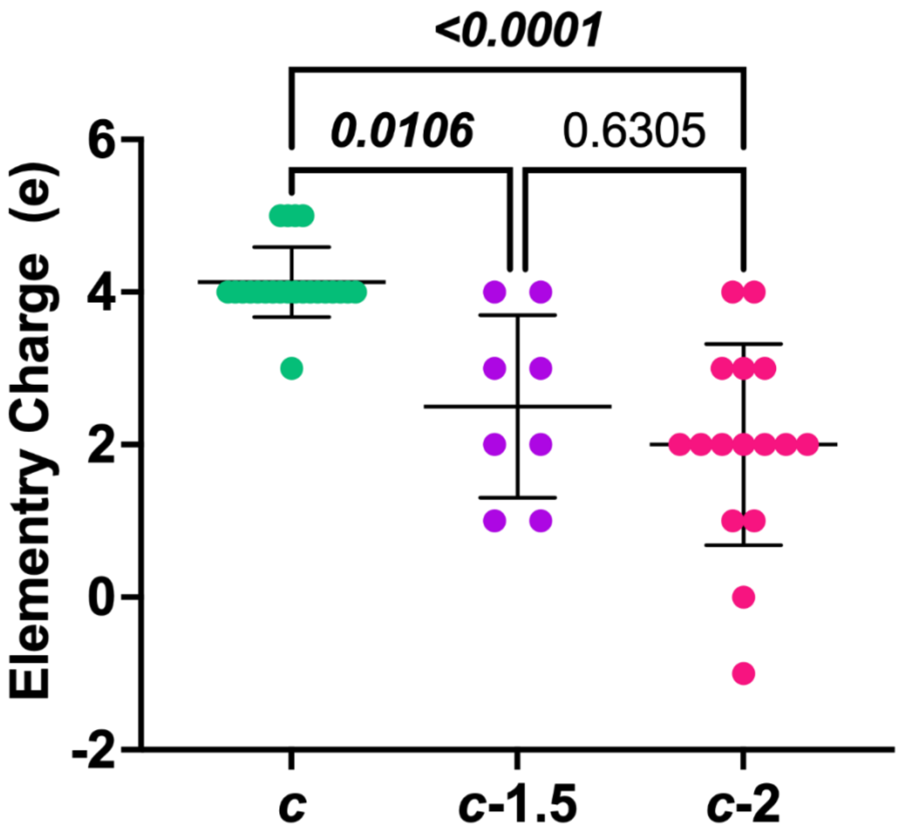


**Figure S14. Net electrostatic charge within 10 Å of the heme iron in cyt c-2 homologs.** Solved crystal structures or AlphaFold3-derived protein/heme c models (from the structural analysis in Fig. S3) were used to quantify the total charge within a 10 Å radius of the heme iron in PyMOL. Statistical differences among groups were assessed using a Kruskal–Wallis test followed by Dunn's post hoc test. The p-values are shown.

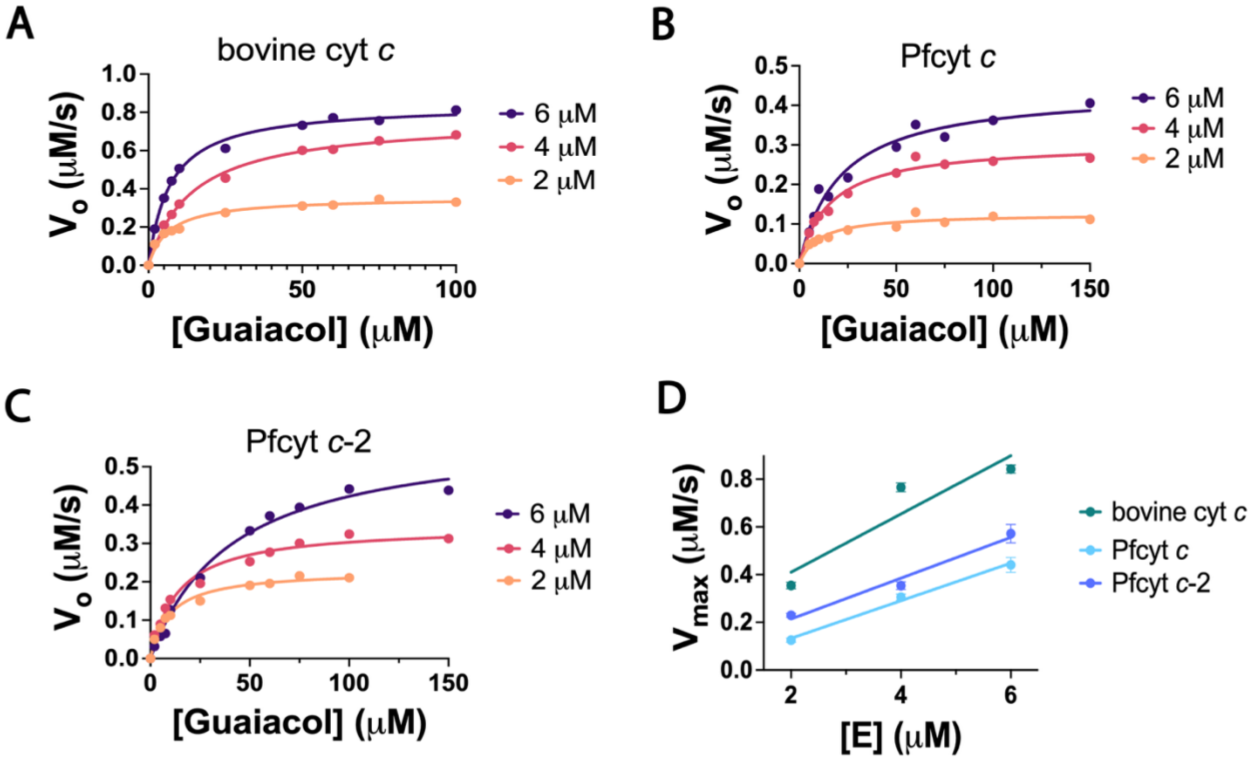


**Figure S15. Michaelis-Menton curves for conversion of guaiacol to tetraguaiacol at three enzyme concentrations**. Activity assays were performed for bovine cyt *c* (A), Pfcyt *c* (B), and Pfcyt *c*-2 (C). (D) $V_{max}$ as a function of enzyme concentration was fit to a simple linear regression that yielded the following $R^2$ values: bovine cyt *c* = 0.86, Pfcyt *c* = 0.99, and Pfcyt *c*-2 = 0.98.

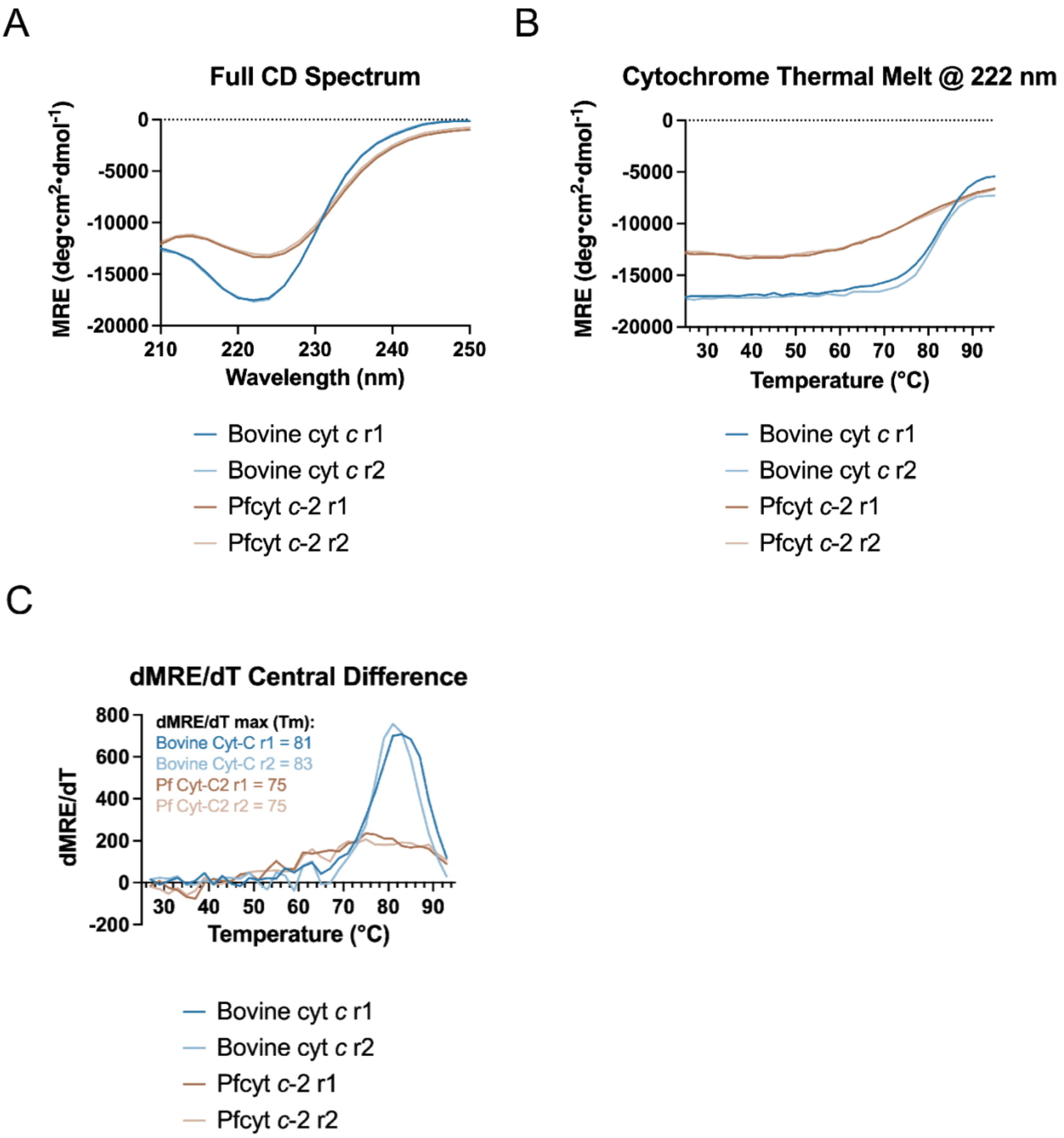


**Figure S16. Circular dichroism (CD) analysis of cytochrome stability.** Data from two independent experiments are shown for bovine cyt *c* (brown) and Pfcyt *c*-2 (blue). (A) CD spectra collected from 210 to 250 nm. (B) Thermal melts monitored at 222 nm. (C) dMRE/dT plots derived from the melt-curve data for each experiment.

### MATERIALS AND METHODS:

**Bioinformatic analysis.** Canonical Pfcyt *c* (Uniprot: Q8IM53) and Pfcyt *c*-2 (Uniprot: Q8I6T6) were used as bait sequences to identify homologs by PSI-BLAST across major eukaryotic clades. Up to ten representative sequences per clade were selected for phylogenetic and sequence similarity network analyses. Protein sequences were aligned using Clustal Omega[7], and the resulting multiple sequence alignment was used as input for phylogenetic inference. A maximum-likelihood phylogeny was generated with IQ-TREE v2[8] using ModelFinder for best-fit substitution model selection and node-support estimation via 1,000 ultrafast bootstrap replicates and 1,000 SH-aLRT replicates. The final tree was visualized using the Interactive Tree of Life (iTOL) webserver[9]. Sequence similarity networks were generated with the EFI-EST webserver and visualized in Cytoscape. Application of a 54% sequence-identity threshold resolved three distinct protein clusters[10].

**Structural Modeling.** Structural models of cytochromes identified in the bioinformatic analysis were generated using the AlphaFold3 server, with heme c included in each prediction[11]. Predicted structures were compared to the solved crystal structure of bovine cytochrome *c* (PDB: 2B4Z)[6] by using PyMOL's align function to calculate the RMSD of the Cα atoms. Coordination bond distances between the iron atom and the axial ligands His19 and Met81 were measured. Electrostatic surface potentials were calculated using the APBS plugin (pH 7.0, 150 mM ionic strength). The net charge within a 10 Å radius of the heme iron was determined by summing the partial charges of all atoms within this sphere.

**Recombinant Protein Expression and Purification.** Bovine cytochrome *c* (>95% purity) was purchased from Sigma (catalog C2037) and used without further purification. For purification of parasite cytochromes, BL21/DE3 *E. coli* were co-transformed with the pET28 plasmid (encoding PfCyt *c*, PfCyt *c*-2, or the mutant PfCyt *c*-2 M81A with N-terminal His-tag) and the pGEX plasmid (encoding the human HCCS with N-terminal GST), as previously reported[1]. Hemylated PfCyt *c* and PfCyt *c*-2 were purified from cell lysates by Ni-NTA affinity chromatography, followed by SP ion-exchange chromatography, and gel filtration chromatography, as previously published[1]. Hemylated PfCyt *c*-2 M81A was purified similarly but without ion-exchange chromatography. Coomassie-stained SDS-PAGE and LC-MS/MS were used to validate the purity and identity of the purified cytochromes, as previously published[1].

**Reduction Potential Determination.** The reduction potentials of hemylated cytochromes were measured using a previously described method[12]. Reactions were prepared in 50 mM potassium phosphate buffer at pH 7.0. Anaerobic conditions were established by adding glucose (5 mM final) and glucose oxidase (50 µg/mL final) and providing xanthine (300 µM final) as an electron source. Once $O_2$-free conditions were achieved, freshly prepared catalase (5 µg/mL final) and purified cytochrome (4 µM final) were added, and the mixture was thoroughly mixed.

A redox dye with a midpoint potential close to that of the cytochrome (e.g., dichlorophenolindophenol (DCIP, 217 mV), Neutral Red (-325 mV), or Safranine T (-289 mV) was then titrated into the solution to produce an absorbance peak similar in height to the cytochrome Soret band. Reduction of the dye–protein mixture was initiated by adding xanthine oxidase (50 nM final). Absorbance spectra from 350–750 nm were recorded every 30 s on a ThermoFisher Evolution 350 spectrophotometer until no further spectral changes were observed (~40 min). Excess sodium dithionite was added at the end of each experiment to obtain fully reduced spectra.

Data were fit to a Nernst plot using wavelengths corresponding to the dye peak and the cytochrome Soret peak. For Neutral Red experiments, the dye showed substantial spectral overlap with the cytochrome Soret region. To account for spectral overlap, stepwise UV–vis spectra of Neutral Red undergoing reduction were collected, and linear regression was used to define the relationship between absorbance at 480 nm and 417 nm. This regression equation was then applied to each Pfcyt *c*-2 titration to estimate the Neutral Red contribution to the Soret peak at 417 nm, which was subsequently subtracted during analysis based on the measured absorbance at 480 nm. All reported potentials (E°) are referenced to the standard hydrogen electrode (SHE).

**Cloning.** The M81A Pfcyt *c*-2 mutant was generated by PCR amplification from WT Pf cyt *c*-2 pET28a plasmid[1] using mutagenic primer sets 1/2 and 3/4 (1: cgcggcagccatatgaat, 2: tttgaagttagctcctatgtgtgactcaacaaattgttgtgg 3: ataggagctaacttcaaaggtttatcgaatttgcaagaaagg 4: gtggtggtggtggtggtg) that introduced the M81A substitution. The amplified product was cloned into pET28a using ligation-independent insertion (NEB E2621) between the NcoI and XhoI restriction sites, positioning the gene in-frame with the vector's N-terminal 6x His tag. The final construct was verified by whole plasmid sequencing (Plasmidsaurus).

**SDS-PAGE and western blot analyses.** For analysis of Pfcyt *c*-2 hemylation in *E. coli*, BL21(DE3) bacteria co-expressing human HCCS-GST (encoded by pGEX vector with N-terminal GST tag) and either wild-type or mutant Pfcyt *c*-2 (encoded by pET28a vector with N-terminal His6 tag) were grown at 37°C in LB media until reaching an optical density at 600 nm of 0.5. Cultures were induced with 1 mM IPTG (GoldBio I2481C50) and 100 µM 5-aminolevulinic acid, then grown overnight at 20°C. Bacteria were harvested by centrifugation, washed, and resuspended in 1× PBS (137 mM NaCl, 2.7 mM KCl, 10 mM $Na_2HPO_4$, 1.8 mM $KH_2PO_4$, pH 7.4) supplemented with 1% Triton X-100 (v/v) and protease inhibitor cocktail (ThermoFisher J61852.XF). Bacteria were lysed by sonication (20 pulses, 50% duty cycle, 50% power) on a Branson sonicator equipped with a microtip, followed by incubation at 4°C on a rocker for 1 hour. Lysates were clarified by centrifugation (10 min at 14,000 rpm). Equal volumes of wild-type and mutant Pfcyt *c*-2 lysates were dissolved in 1× SDS sample buffer, heated at 95°C for 10 minutes, and fractionated by SDS-PAGE using 12.5% acrylamide gels run at 120 V in the Bio-Rad mini-PROTEAN electrophoresis system. Proteins were transferred to nitrocellulose membrane for 1 hour at 100 V using the Bio-Rad wet-transfer system. For evaluation of hemylation, membranes were developed for 5 min with Prometheus ProSignal Femto ECL reagent (Genesee Scientific 20-302) and imaged using an Invitrogen iBright 1500 Imaging System. Membranes were then stripped and reprobed for evaluation of cytochrome *c* and HCCS protein expression via western blot. Membranes were blocked with 5% (w/v) skim milk in 1× PBS for 1 hour at ambient temperature, probed at 1:1000 dilution with mouse monoclonal anti-GST-DyLight800 conjugated antibody (ThermoFisher MA4-004-D800) and mouse monoclonal anti-His6 tag-DyLight680 conjugated antibody (ThermoFisher MA1-21315-D680), washed in 1× PBS supplemented with 0.5% (v/v) Tween-20, and imaged on a LI-COR Odyssey CLx.

**Circular dichroism.** All circular dichroism (CD) studies were completed on an Aviv Biomedical Inc. Model 410 CD spectrometer using a 1 cm quartz cuvette with a stir bar spinning at 30%. All samples were run at 2 μM[13] in PBS supplemented with 10 μM ammonium persulfate (to ensure oxidized protein state). Spectra for each sample were measured in triplicate from 250-210 nm, measuring every 2 nm with 10 seconds of averaging time at each point. Thermal melt studies measured CD signal at 222 nm, climbing every 2 °C from 25-95 °C. Each temperature was equilibrated for 1 minute before data collection over a 10 second averaging time. The first derivative of the thermal melt curve was calculated using the central difference method (algebraic calculation of slope using the point before and point after each temperature). $T_m$ was calculated as the temperature at the maximum of each first derivative spectrum.

**Guaiacol Assay of Peroxidase Activity.** Guaiacol assays were performed at 25 °C by monitoring the formation of tetraguaiacol at 470 nm under previously reported conditions[4, 14]. Stock solutions of $H_2O_2$ (100 mM, Fisher BP2633) and guaiacol (500 μM, Sigma Aldrich G5502) were verified spectrophotometrically using extinction coefficients of $\varepsilon_{240}$ = 41.5 $M^{-1}$ $cm^{-1}$ for $H_2O_2$ and $\varepsilon_{274}$ = 2150 $M^{-1}$ $cm^{-1}$ for guaiacol[14]. A 2× mixture containing purified cytochrome and varying concentrations of guaiacol was prepared in 50 mM potassium phosphate buffer, pH 7.0. This mixture was combined 1:1 with a 100 mM $H_2O_2$ stock solution to yield final concentrations of cytochrome (2, 4, or 6 μM), guaiacol (0, 2, 5, 7.5, 10, 25, 50, 60, 75, or 100 μM), and 50 mM $H_2O_2$. Immediately upon addition of $H_2O_2$, absorbance at 470 nm was recorded using a ThermoFisher Evolution 350 spectrophotometer until tetraguaiacol formation reached a plateau. Rates of guaiacol consumption (μM $s^{-1}$) were calculated from the change in absorbance at 470 nm over the initial linear phase using the extinction coefficient for tetraguaiacol ($\varepsilon_{470}$ = 26.6 $mM^{-1}$ $cm^{-1}$)[4]. Average initial rates from three replicates at each guaiacol concentration were fit to Michaelis–Menten curves in GraphPad Prism 10.6 to determine $k_{cat}$ and $K_m$.
